# Nascent clathrin lattices spontaneously disassemble without sufficient adaptor proteins

**DOI:** 10.1101/2021.04.19.440502

**Authors:** Si-Kao Guo, Alexander J. Sodt, Margaret E. Johnson

## Abstract

Clathrin-coated structures must assemble on cell membranes to perform their primary function of receptor internalization. These structures show marked plasticity and instability, but what conditions are necessary to stabilize against disassembly have not been quantified. Recent *in vitro* fluorescence experiments have measured kinetics of stable clathrin assembly on membranes as controlled by key adaptor proteins like AP-2. Here, we combine this experimental data with microscopic reaction-diffusion simulations and theory to quantify mechanisms of stable vs unstable clathrin assembly on membranes. Both adaptor binding and dimensional reduction on the 2D surface are necessary to reproduce the cooperative kinetics of assembly. By applying our model to more physiologic-like conditions, where the stoichiometry and volume to area ratio are significantly lower than *in vitro*, we show that the critical nucleus contains ^~^25 clathrin, remarkably similar to sizes of abortive structures observed *in vivo*. Stable nucleation requires a stoichiometry of adaptor to clathrin that exceeds 1:1, meaning that AP-2 on its own has too few copies to nucleate lattices. Increasing adaptor concentration increases lattice sizes and nucleation speeds. For curved clathrin cages, we quantify both the cost of bending the membrane and the stabilization required to nucleate cages in solution. We find the energetics are comparable, suggesting that curving the lattice could offset the bending energy cost. Our model predicts how adaptor density controls stabilization of clathrin-coated structures against spontaneous disassembly, and shows remodeling and disassembly does not require ATPases, which is a critical advance towards predicting control of productive vesicle formation.

**Significance Statement:** Stochastic self-assembly of clathrin-coated structures on the plasma membrane is essential for transport into cells. We show here that even with abundant clathrin available, robust nucleation and growth into stable structures on membranes is not possible without sufficient adaptor proteins. Our results thus provide quantitative justification for why structures observed to form *in vivo* can still spontaneously disassemble over many seconds. The ATPases that drive clathrin disassembly after productive vesicle formation are therefore not necessary to control remodeling during growth. With parameterization against *in vitro* kinetics of assembly on membranes, our reaction-diffusion model provides a powerful and extensible tool for establishing determinants of productive assembly in cells.

## INTRODUCTION

In cells, clathrin-coated structures assemble into membrane bound puncta^1^ that proceed to productive vesicles only about half the time^2^, otherwise, they disassemble. These structures are observed to contain at least ^~^20 clathrin^1^; are they dynamic^3^ and unstable? Many proteins have been shown to tune the frequency and probability of disassembly in these transition structures ^1,2,4,5^, but none of these proteins are physical drivers of disassembly. The clathrin *uncoating* machinery, which is capable of driving disassembly^6,7^, is rarely seen at maturing clathrin-coated structures^8,9^. A fundamental question thus remains, what physically stabilizes clathrin-coated structures against disassembly, and to what extent? Addressing this question will help to establish under what conditions early clathrin-coated structures develop into productive vesicles. Here, our simulations of clathrin recruitment and assembly on membranes reproduce *in vitro* kinetic data, validating a model that then establishes how the key parameters of adaptor density, volume to area (V/A) ratio, and clathrin concentration determine the kinetics and critical nuclei of clathrin-coated structures on membranes.

Experiments *in vitro* have shown that clathrin-coated structures can assemble robustly on membranes with clathrin and a minimal set of components, requiring a cytosolic adaptor that localizes clathrin to the membrane and membrane binding sites that localize adaptors to the membrane, which physiologically is the essential lipid PI(4,5)P_2_ ^10–12^. In addition, the *in vitro* clathrin assembles both flat and vesicle-shaped lattices, just as is observed *in vivo*^13,14^. What these experiments have not probed, however, is what stoichiometry of clathrin, adaptors, and membrane area determines the transition from unstable to stable clathrin coats, as all conditions studied report equilibrium, highly stabilized lattices. Even when component concentrations are comparable to *in vivo* reported values, the V/A ratio is typically orders of magnitude higher than in cells, which can dramatically increase stability of membrane associated complexes^15^. *In vivo* experiments have provided more detailed insight into the dynamics of clathrin coat nucleation and budding^3,16^, but the number of factors that are known to contribute to clathrin-mediated endocytosis in cells, including enzymatic activity^17,18^, makes it impossible to estimate the critical nucleus of clathrin and adaptors that is stabilized against disassembly. The composition of successful productive vesicles^19^ demonstrates that a diversity of adaptor and receptor compositions result in productive budding, and the goal of our study is to establish minimal criteria for stable nucleation and growth dependent on adaptor concentration, out of equilibrium.

Kinetic measurements have provided a powerful variable to assess both models and mechanisms of solution assembly in diverse filament forming^20,21^, aggregating^22,23^, and capsid forming systems^24^. With recent kinetic experiments, Sarkar and Pucaydil^25^ provided insight not only on the initial conditions that can drive stable clathrin lattice formation on membranes, but constraints on rates of growth, while previous biochemical experiments constrain the relative free energy of clathrin-clathrin^26^ and clathrin-adaptor^27^ interactions. The kinetic experiments used initial conditions that again drove stable lattices, and could not test the requirements for stable vs unstable lattices. To assess the size and speed of stable nuclei as they might occur in the cell, we need to mimic the lower V/A ratio of the cell, recreating these experiments *in silico* at new conditions that are challenging experimentally. A great advantage of the type of microscopic spatial modeling here is that instead of requiring experiments at many distinct concentrations in order to fit rate equations to the data^23^, our explicit physical models tightly constrain the subset of rate constants that can quantify the data.

Modeling has played an important role in understanding principles of clathrin-coated assembly, but has yet to characterize the kinetics of assembly, due to the challenges of capturing molecular structure at mesoscopic length (μm) and time (minutes) scales. We achieve this here using recently developed structure-resolved reaction-diffusion software^28^, which captures coarse molecular structure^29^ for multicomponent systems in 3D^30^, 2D^31^, and transitioning between^32^. Critically, dimensional reduction, or the change in search space and dynamics that accompanies transitions from 3D to 2D^33^, is rigorously accounted for in our model, which will quantitatively impact stability^15^ and kinetics^34^ of assembly steps. Our model produces energetics of clathrin cages similar to purely statistical mechanical models ^35–39^, although those models lack temporal resolution and molecular detail. Our model produces similar structures to previous spatial simulations in solution^40–42^, but with membrane included these simulations were too expensive to characterize experimental kinetics^43^. We here estimate the energetic balance of curved cage formation and membrane bending by determining the minimum energy membrane shape compatible with curved solution cages. Similarly to other approaches^44–46^, we do not dynamically couple cage assembly to vesicle formation. Yet we are uniquely able to capture kinetics of localization and assembly on membrane by tracking flat lattice assembly, which is consistent with observed early growth ^13^. Our model provides a starting point to move beyond non-spatial kinetic models^47,48^ or force-balanced models^49^, and build in essential structural details that will necessarily impact the packing, mechanics, and remodeling of adaptors and cargo in vesicles. Critically, our simulations provide details on each clathrin monomer and higher-order structure as they evolve in time and space. Using concepts from classical nucleation theory, we can thus quantify the critical nuclei as a function of adaptor density, showing that at fixed clathrin concentration, a minimum adaptor density is necessary to drive nucleation and continued growth at relevant physiologic speeds (^~^60s).

## MODEL

Our model is designed to reproduce the quantitative kinetics of recently measured clathrin assembly on membranes^25^ by capturing the coarse structure of the clathrin trimer and its adaptor protein, protein (and multi-protein) diffusion constants, rates describing all pairwise protein-protein interactions, and the concentrations and V/A ratio of the experiment. Each rigid clathrin trimer contains 3 sites for binding other clathrin, arranged to produce a flat or curved hexagonal lattice depending on the ‘pucker’ angle *α* of the legs relative to the z-axis (Fig. 1). Each trimer also contains 3 sites to bind adaptors. Each adaptor contains a single site for clathrin, and a single site to bind a membrane lipid. When two molecules bind via their specific interaction sites, they adopt a pre-specified orientation relative to one another, producing either a flat hexagonal lattice (*α*=90°), or a spherical cage (*α*>90°), with the cage composed of hexagons, pentagons and imperfect contacts due to the rigidity of the trimers^28^. The lengthscales of the proteins are chosen to match the known size of the clathrin lattice^50^ with a center-to-center distance of 17nm here. Excluded volume (Fig 1) is essential to prevent lattices from overlapping as they diffuse in 3D or 2D, which we verified did not alter the kinetics (Fig. S1A).

**Figure 1.**
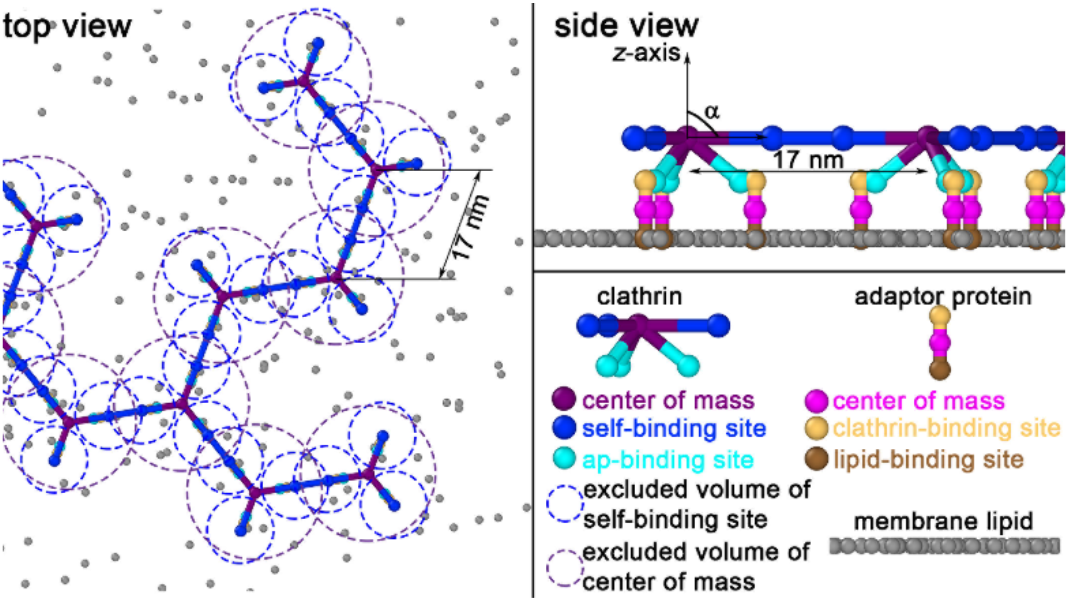
The structure-resolved reaction-diffusion model for clathrin recruitment and assembly on membranes captures structure, valency, and excluded volume. Clathrin trimers are initialized in solution, but bind to adaptor proteins that are localized to the surface. The binding of clathrin to adaptors only once on the surface is consistent with the behavior of AP-2^10^. Adaptor proteins can bind specific lipids on the membrane surface if initialized in solution. Unbound clathrin-clathrin sites exclude one another with a radius of σ=5nm (blue circles), and clathrin-adaptors with σ=1nm. At all times in the simulation, center-of-mass (COM) sites on trimers exclude other trimer COMs at σ =10nm (purple circles).

We fixed several parameters based on experiment, described here. We use fixed dissociation constants of 120μM for clathrin-clathrin (without adaptor)^26^ and 25μM for clathrin-adaptor^27^. The experimental V/A ratio is 991μm (Methods), with a clathrin concentration of 80nM^25^. Our simulation volume has a flat SA of 1μm^2^, which would require a height of 991μm. To retain this same V/A ratio without propagating a huge excess of solution clathrin, we instead use a height of 1μm and maintain a constant (stochastically fluctuating) concentration of [Cla]_tot_=80nM (48 copies) in solution (Fig. 2A and see SI Methods). This assumes the total concentration of clathrin in solution does not change as clathrin accumulates on the membrane, which is true to a very good approximation, with <5% of total clathrin ending up on the membrane by 100s. For the kinetic simulations, our membrane is modeled as a flat surface, and individual binding sites on the surface are modeled implicitly^32^. The implicit adaptors have a height of 4nm, which ensures accurate membrane binding kinetics, as verified by changing the simulation time-step (Fig. S1B) and by a simple system without the clathrin-clathrin reaction (Fig. S1C). We estimate transport properties from Einstein-Stokes, with D_Cla_=13μm^2^/s, D_R,Cla_=0.03 rad^2^/s, D_ap_=25μm^2^/s, D_R,ap_=0.5 rad^2^/s, D_lipid_ = 0.5μm^2^/s, and D_R,lipis_=0.01rad^2^/s, to allow bound complexes to rotate on the surface. Diffusion slows as complexes grow consistent with Einstein-Stokes^29^.

**Figure 2.**
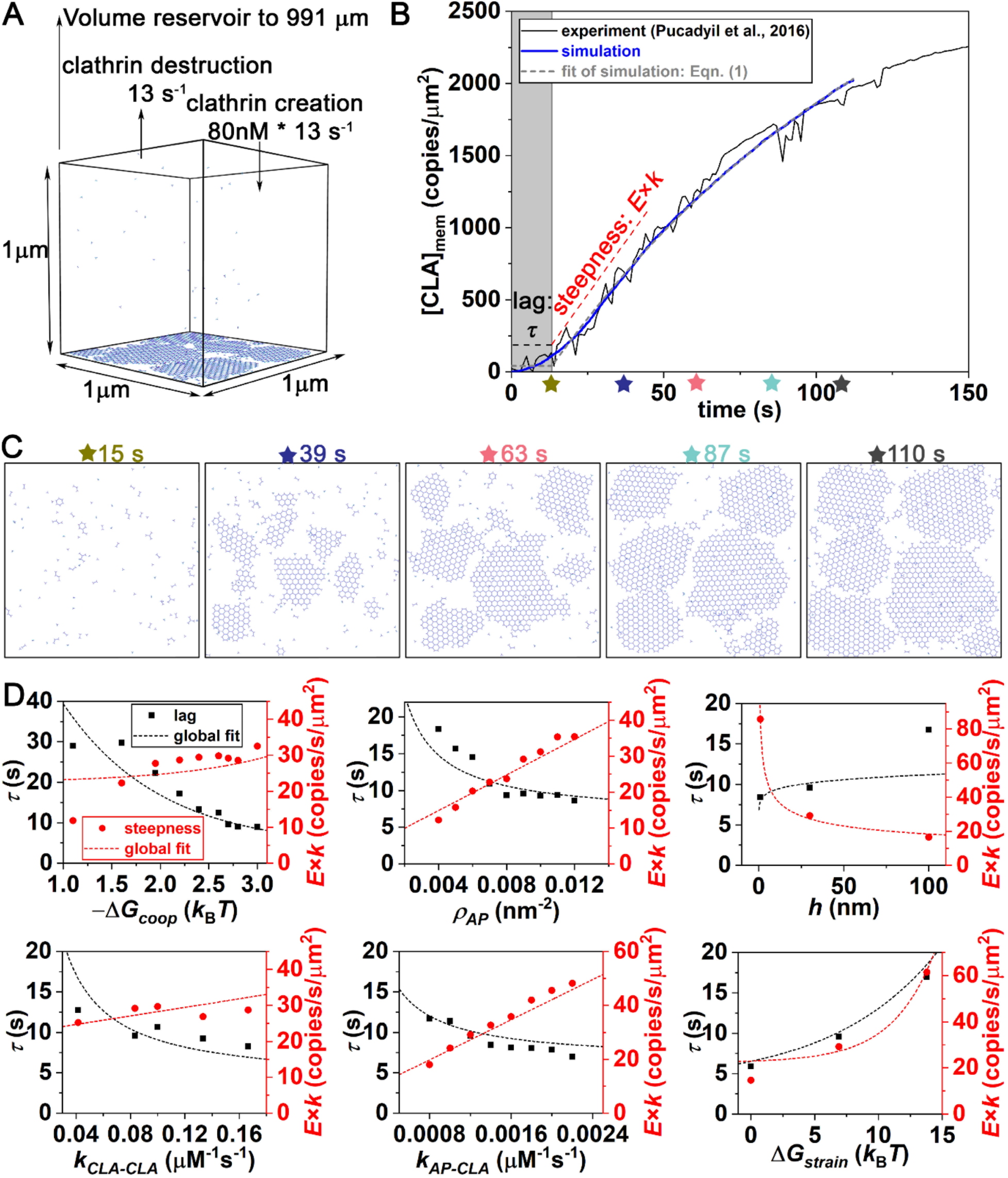
Structure-resolved reaction-diffusion simulations of clathrin assembly kinetics on membranes reproduce *in vitro* experiment. A) Simulations were set up to mimic experimental conditions from published work of Pucadyil *et al* ^25^, where fluorescence of clathrin on membrane tubules was measured with time. A constant (fluctuating) solution concentration of clathrin (at 80nM) was maintained through exchange with the large volume reservoir. B) Fluorescence data averaged over multiple tubules is shown in black. The experimental data was plotted in units of copy numbers per μm^2^, not arbitrary units, by zeroing the offset (subtracted off 300) and rescaling the height by 1.7. The model result is shown in blue (averaged from 4 simulation trajectories). C) Snapshots of one simulation trajectory at different time points. D) Macroscopic time-scales (lag time *τ* and initial growth rate E*k—see Eq 1) as we vary model parameters of Table 1, along with fits (Eq 2 and 3).

The parameters that we varied to optimally reproduce experiment are summarized in Table 1. We allow the relative on and off rates of clathrin-clathrin (*k_CLA-CLA_*)and clathrin-adaptor (*k_AP-CLA_*) binding to vary. Our clathrin-clathrin K_D_ is strengthened when a clathrin is bound to an adaptor, which has been long known from experiment^51^. We thus lower the free energy, Δ*G*_+*ap*2_ = Δ*G*_*coop*_ + Δ*G_*coop*_* using Δ*G_coop_* values in a range −1.1 to −3*k_B_T*, where *k_B_* is Boltzmann’s constant and T is the temperature. We keep the clathrin-clathrin off-rate unchanged and accelerate the on-rate by *f_coop_* = exp (–Δ*G_coop_/k_B_T*). Our model accounts for the changes to rates that accompany localization to the 2D membrane^52^, which is compactly defined by a single molecular length-scale, *h*. We tested values of *h*=1-100nm, the order-of-magnitude of the molecular clathrin length-scale. We introduce a strain energy Δ*G_strain_* for the formation of closed polygons (hexagons or pentagons). When the clathrin lattice forms a closed hexagon (or pentagon), dissociation of a single clathrin-clathrin bond does not release a monomer or fragment. The ratio of rebinding to unbinding rate is given by 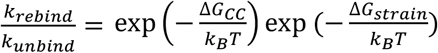, where Δ*G_CC_* is the free energy difference of an unbound to bound pair of clathrin^28^. The term exp 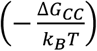 is simply 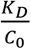 for the pairwise reaction, where *C*_0_ is the standard state concentration (1M), giving a rebinding ratio of 8.3×10^3^ or 9.2×10^4^ for clathrin-clathrin bonds without and with adaptor bound, respectively. The much faster rebinding rates render hexagons highly stable against dissociation. The term exp 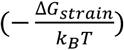 thus lowers the stability of a hexagon compared to 6 ideal bonds (for Δ*G_strain_* > 0). There are several reasons we would expect this because clathrin is a flexible molecule that forms varied lattices and cages. Forming two bonds to close a hexagon/pentagon could place elastic strain on the clathrin, reducing its stability. Some *in vivo* lattices, particularly the flat lattices, also form highly imperfect ‘hexagons’ that clearly have poorly aligned contacts^53^, contributing to a less stable and more dynamic lattice. With a Δ*G_strain_* value of +6.9*k_B_T*, the free energy of the hexagonal 6-mer corresponds to the strength of 5.4 ideal bonds rather than 6, whereas a 6-mer in an extended conformation contains 5 bonds. Lastly, the density of adaptors (*ρ_AP_*) in the *in vitro* simulations are treated as a variable parameter because although the density of membrane binding sites was known (0.0746/nm^2^), we would not expect all these sites to recruit adaptors. For a K_D_ ranging from 1-10μM ^54^ between chelator and His-tagged adaptor (present at 200nM)^25^, we would have ^~^0.0014-0.012/nm^2^ adaptors on the surface before the clathrin is flowed in, which we use as limits on our adaptor density parameter. We note that in the *in vitro* experiments, only the beta appendage of AP-2 is present, and we assume that it remains bound to the membrane throughout the simulations. In the ‘physiologic-like’ conditions, we allow the AP-2 protein to reversibly bind the membrane, and prevent this full-length AP-2 from binding to clathrin in solution because binding is inhibited prior to membrane localization^10^.

**Table 1.**
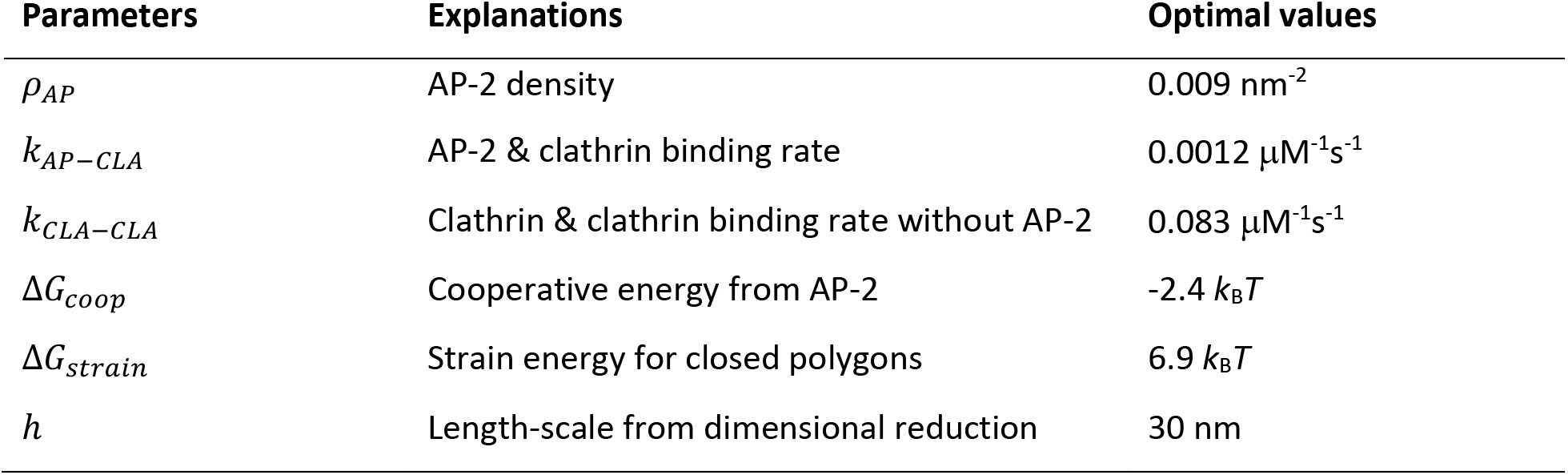
The varying parameters in the model and optimal values to reproduce the experimental kinetics

## RESULTS

### Quantitative agreement established between simulation and *in vitro* kinetics

Simulations of clathrin lattice growth (see Fig. 2A) reproduce experimental kinetics under equivalent conditions (Fig. 2B). The primary features of the kinetics are the lag time (*τ*) and the slope of the initial growth (*k* × *E*), which follow from an approximate expression for membrane-bound clathrin, [CLA]_mem_(*t*):

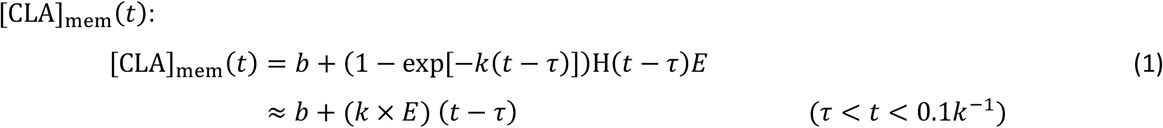

Mathematically, the lag in initial growth is accounted for by the Heaviside step function H, which is zero before *τ* and one thereafter. Simulated timescales [*τ*, 1/*k*] = [13s,121s] are in excellent agreement with those of experiment [11s,107s]. The geometry of the experiment (*V/A*) and size of the clathrin lattice limit [CLA]_mem_ to ^~^5000 clathrin/um^2^ (^~^201nm^2^ per trimer). In both experiment and simulation, saturation of clathrin on the surface thus results from a loss of accessible surface area rather than a loss of adaptor binding sites. Rapid growth following an initial lag is a hallmark of cooperativity. Here, clathrin binds adaptors, increasing its association rate with other clathrin. The reduced geometric search space following membrane binding (dimensional reduction) further increases the rate.

### Cooperativity from both dimensional reduction and adaptor binding are required to reproduce experimental kinetics

Experiments have established that once clathrin binds the adaptor proteins, its affinity for other clathrins is increased^51^. Based on our simulations, where the K_D_ without adaptor is fixed^26^, we observe that the affinity with adaptor is increased by ^~^2.4 *k_B_T*, leading to faster and more stable binding. Further, once clathrin is localized to the surface, it binds additional adaptors or clathrins with a 2D rate constant^15^, with a length-scale of 30nm determined by comparison with experimental kinetics (see Modeling), that is comparable to clathrin’s radius. Both sources of cooperativity contribute significantly to rapid growth on the surface (Fig. 2D).

### Lag time is controlled by the sticking rate of clathrin to the surface

Figure 2D reports, in black, the sensitivity of *τ_lag_* to six principal parameters of the model (see Table 1). The simulation data shows clear trends that are well fit by a simplified model summed over two timescales, the first representing clathrin localization to the membrane, the second nucleation of clathrin-clathrin contacts:

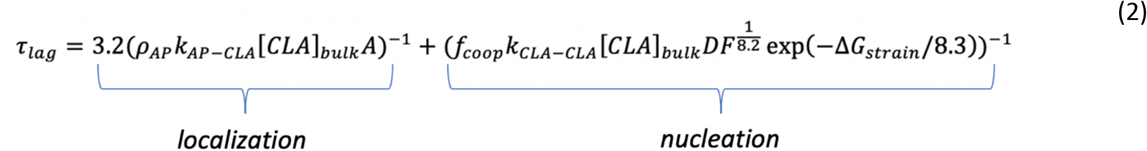

where *f_coop_* = exp (–Δ*G_coop_/k_B_T*) and the dimensionality factor *DF = V/A/h*. [*CLA*]_*bulk*_ is the clathrin concentration in the solution. The simplified model of *τ_lag_* is plotted as a black dashed line in Fig. 2D. The parameters of the simplified model are determined by fitting all simulations results (including Figs. 2–3), so that it describes both *in vitro* and the physiologic-like simulations below (Fig. S2, SI Methods, Table S2). The dependence of *τ_lag_* on the rates and concentrations are similar to a half-time^55^ or mean first passage time^34^, varying inversely with *rate* x *concentration* in both terms. However, there is a nonlinear dependence on the dimensionality factor (*DF*) and the elastic strain of the clathrin lattice (Δ*G_strain_*). This is likely because some clathrins bind directly to clathrin from solution, skipping the cooperative mechanisms provided by dimensional reduction. Nucleation at the surface is also less dependent on the total volume, as it siphons only a small number of clathrin. In these *in vitro* simulations, the localization time of clathrin to the membrane contributes about 40% of the lag (first term in Eq 2), and is, as expected, most sensitive to both the rate of clathrin binding to adaptor and the density of adaptor on the membrane. Nucleation of a clathrin structure (second term) requires multiple clathrin to localize at one position, which is limited primarily by the low concentration of clathrin (0.08μM).

**Figure 3.**
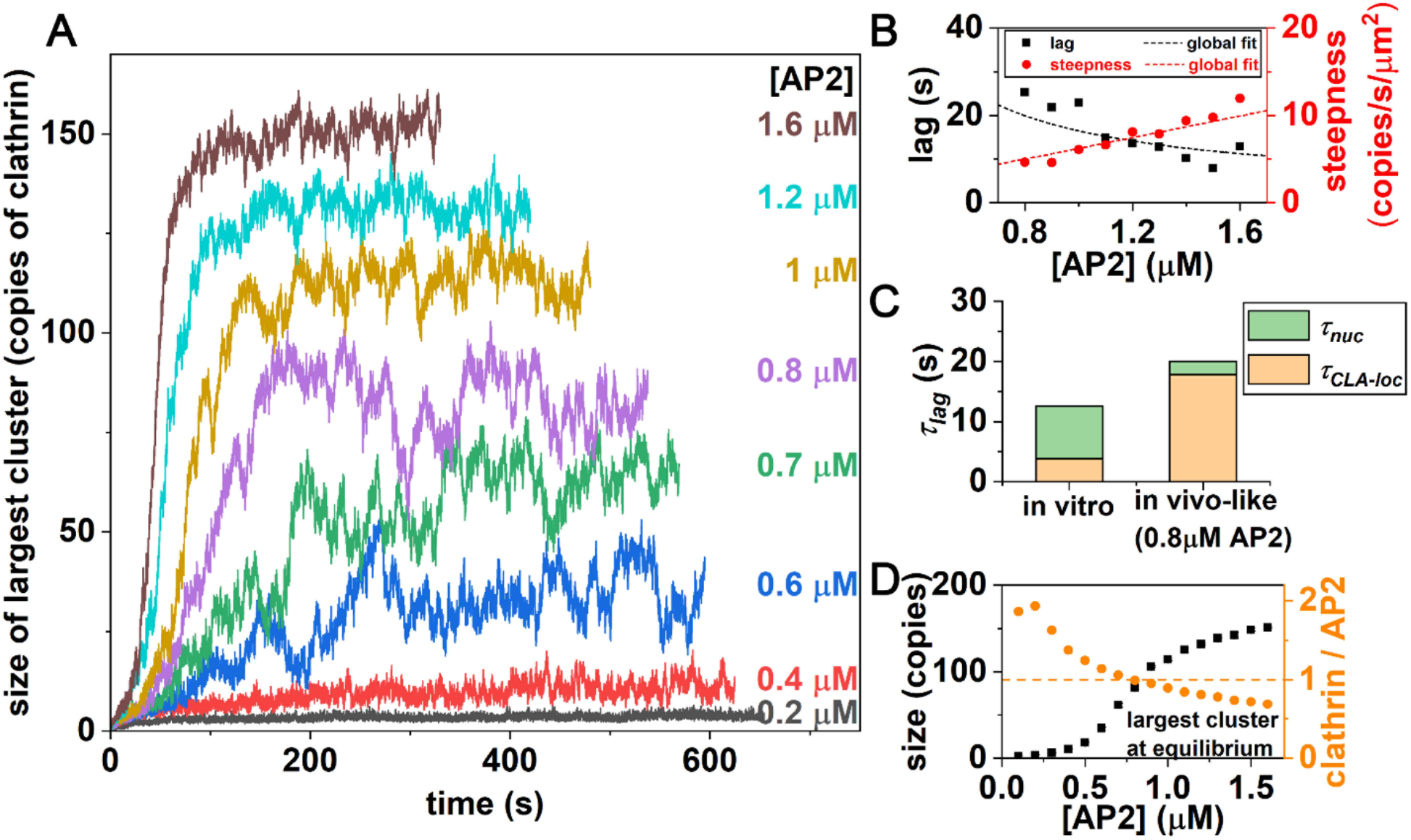
Clathrin lattices at physiologic-like geometries fail to assemble until adaptor concentration reaches 0.6μM. A) Kinetics of largest clathrin lattice assembly is faster and reaches larger sizes with increasing adaptor concentration. All simulations have 0.65μM clathrin. B) Lag times and growth rates follow the simple formulas of Eq 2 and Eq 3. C) Lag times are slower in these geometrically physiologiclike simulations and rate-limited by recruitment of clathrin to a membrane with far fewer adaptor binding sites than *in vitro*. D) The size of the largest lattices increases with adaptors, transitioning from small to large lattices at 0.8μM (black squares). At this point, the stoichiometry of clathrin:adaptor has reached 1:1 (orange circles).

### Growth kinetics is controlled primarily by recruitment from adaptors

Figure 2D shows, in red, the sensitivity of the initial growth rate *k* × *E* to the parameters in Table 1. As with *τ*, a simplified model, motivated by mass-action kinetics, fits the data reasonably well in most cases (Fig. 2D, red dashed line):

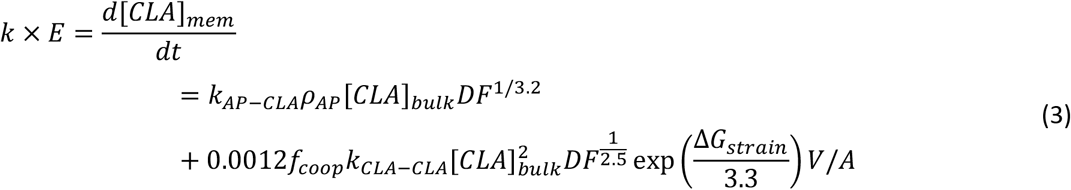

This initial rapid growth is most sensitive to the first term (proportional to *k_AP-CLA_*) accounting for recruitment of clathrin to adaptors. Growth is further enhanced cooperatively due to 2D interactions between both clathrin trimer-trimer and clathrin-adaptor contacts. Varying *h*=*k*_on,3D_/*k*_on,2D_ at fixed V/A shows the sensitivity to the DF (Fig 2D), indicating that 2D binding is necessary to drive experimental growth speeds.

Counter-intuitively, a higher strain free energy Δ*G_strain_* slows the lag time but *increases* the growth rate, whereas all other parameters cause correlated slow-downs in lag and growth. Increasing the strain penalty destabilizes closed hexagonal structures, promoting disassembly which slows nucleation (τ*_lag_*), as we would expect. However, the growth is accelerated, due to increased sampling of dynamic and disordered structures with more ‘sticky ends’ or free legs (1 free leg/trimer in a hexagon, 1.33 free leg/trimer in extended 6-mer), which recruits additional clathrin more rapidly.

### AP-2 alone is insufficient to nucleate lattices at cellular conditions

To characterize nucleation at more physiologic-like conditions, we reduce the *V/A* ratio from 991μm, to 1μm, representative of a HeLa cell, and increase the clathrin trimer concentrations to 0.65μM^56^ (Methods). Stable lattices do not nucleate at physiologic AP-2 concentrations (0.2μM^56^, see Fig. 3A). Compared to the *in vitro* system, the significantly reduced *V/A* results in a much lower density of AP-2 on the membrane (^~^120μm^-2^, compared to 9000μm^-2^). Most clathrin assemblies are dimers and monomers, with a stoichiometry of 2 clathrin:1 AP-2 that is too low to induce the cooperativity necessary to nucleate. At this clathrin concentration, the hallmark lag and growth phases require a clathrin to adaptor ratio of ^~^1:1 (reached at 0.7-0.8μM of adaptor). Equilibrium fluctuations are largest at this point, indicating a transition in lattice growth (Fig. S3). Increased AP-2 nucleates larger lattices, with both shorter lag times and faster growth (Fig. 3B).

Equation 2 indicates the lag time increases significantly at low concentrations of AP-2. With few AP-2 binding sites, localization of clathrin to the membrane is rate-limiting (Fig 3C). The growth rate is significantly slower than *in vitro*, where only a few clathrin are recruited to a μm^2^ surface area per second (Fig 3B). Here again the few AP-2 available limits recruitment of clathrin to the surface. The recruitment of clathrin by clathrin has also slowed, due to the sensitivity of the growth rate to the *DF* in stabilizing clathrin-clathrin contacts on the surface. We note that the recruitment of AP-2 to lipids is not rate-limiting in these simulations, as PI(4,5)P_2_ coverage is ^~^20000μm^-2^ (1%), and the rate of AP-2 to lipid binding is fast, 0.3s^-1^μM^-1^ (Table S2).

### Assembly of lattices overcomes an initial barrier, followed by small increases in stability during growth

Tracking the lattice size throughout a simulation allows quantification of their relative probability during growth in *kT* units, — ln *p*_obs_(*n; t*), (Fig. 4A), and the equilibrium free energy of all lattice sizes, — ln *p*(*n*). At equilibrium, and for higher adaptor concentrations we observe a bimodal distribution—either very large lattices or a population of small lattices of n<^~^25 (Fig. 4A,C). This is a notable outcome, as it demonstrates that even at higher adaptor concentration, after the majority of solution clathrin and adaptor are concentrated into a single coated structure, the remaining clathrin forms small clusters that are not stabilized against disassembly. Hence the formation of stable clathrin lattices can locally deplete resources needed for nucleation of additional stable sites. The growth phase of assembly (Fig. 4A-yellow) further shows how even after overcoming the initial barrier to nucleation, lattices exhibit similar stability at a broad range of sizes, indicating dynamic remodeling (Fig. 4A, 4B). This flat region is followed by a stable well that shifts larger and deepens as the system equilibrates to the maximal lattice sizes, with a final ‘wall’ that prevents further growth. This wall results from insufficient adaptors to stabilize growth at the peripheries of the lattice.

**Figure 4.**
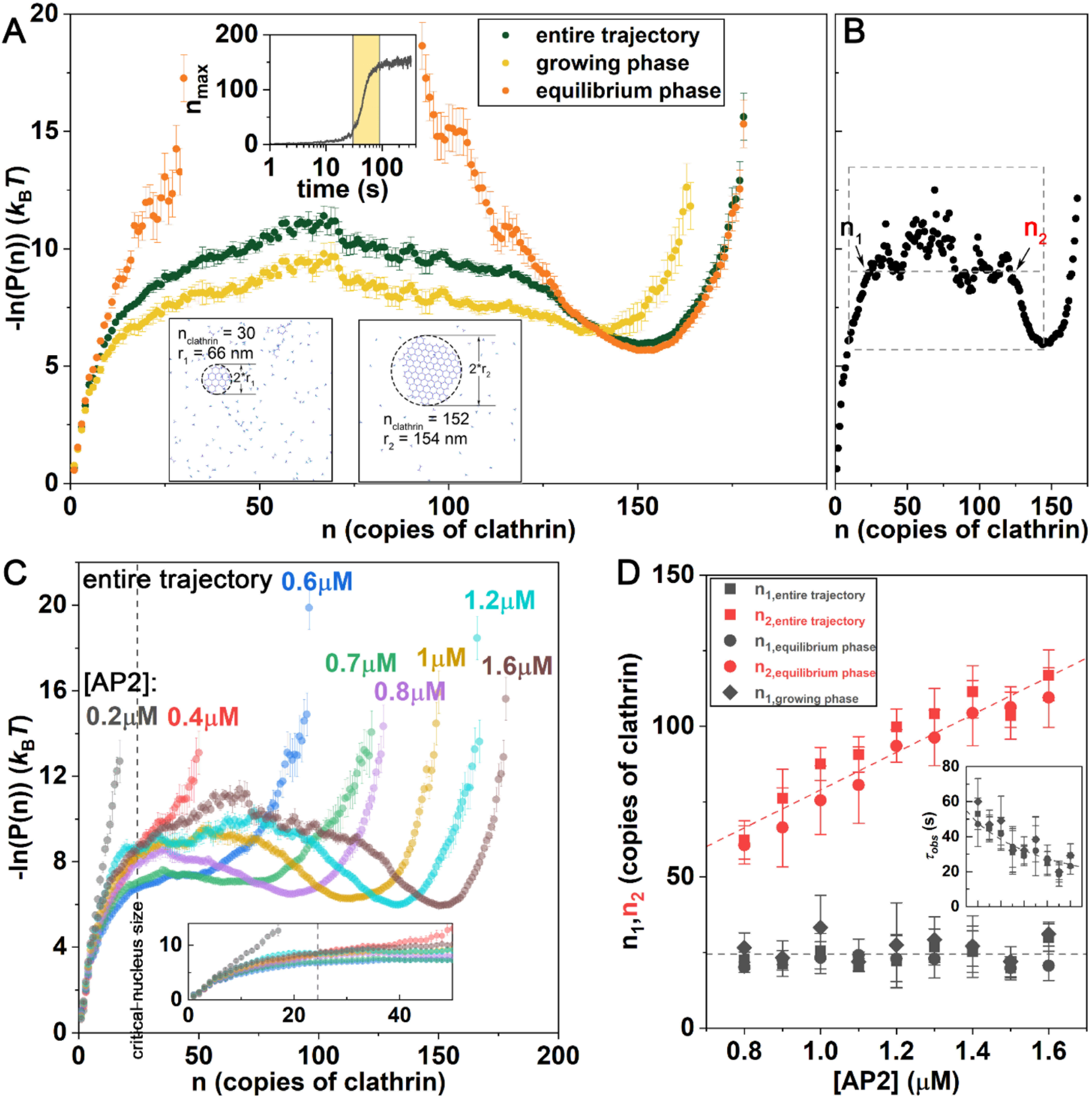
Clathrin lattices face an initial barrier to growth, with a stable size reached only after significant growth. A) The probability of observing clathrin lattices of size *n* can be converted to an energy-like metric -ln(P(*n*)), which at equilibrium is a true free energy. An initial barrier to nucleation plateaus, followed by a flat region, where structures have comparable probability. Growth phase (yellow) samples intermediate size lattices, while at equilibrium only small and large clusters are visible. Full trajectory is in green. [AP2]=1.6μM. B) To quantify the end of the initial barrier to growth (n_1_), and the start to the stabilized growth (n_2_), we define a plateau at constant -ln(P(n)) that defines these intercepts for each trajectory (Methods). C) With increasing adaptor concentration, larger lattices become stabilized. D) The critical size where the barrier plateaus is ^~^n_1_=25, independent of adaptor concentration. The plateau region is followed by a well that starts at n_2_ and increases with adaptors. (Inset) The time to pass the first barrier follows with the inverse of adaptor concentration *τ_obs_* ∝ 1/[*AP2*], with a proportionality constant 1/[0.026*μ*M^-1^s^-1^].

### Adaptor concentration controls the time to reach the observed nucleus, but not its size

As we increase adaptor concentration beyond 0.7μM, we observe that across the full trajectory, the initial barrier to nucleation transitions to a flattened region at a similar lattice size of n_1_^~^25 clathrin for all adaptor concentrations (Fig. 4C–4D). This initial barrier is significant because smaller lattices are prone to disassembly (Fig S3C). There is a later transition where the stable well starts to form, at n_2_, which clearly shifts to larger lattices as the adaptor concentration increases (Fig. 4C). These two lattice sizes, which bound the flattened region that occurs during growth, are quite consistent with the lattices sizes that define the equilibrium regions (Fig. 4D)(Methods). In all phases, there thus remains a barrier to nucleation that is largely independent of adaptor concentration, whereas the maximum size of lattices grows. We do observe some features in the flatter, noisier intermediate region across the full trajectory, where a small barrier can exist at higher lattice size (n^~^40-70). This indicates that during growth, lattices that are about halfway to the stable size are the least commonly observed, and indeed have a short lifetime (Fig S3). Additional adaptors also dramatically accelerate the time required to reach the ‘critical’ nucleus, with a time approximately proportional to inverse adaptor concentration (Fig. 4D-inset).

The size of our observed ‘critical’ nuclei shows that even after multiple (8-10) hexagons (Fig. 4A inset) have formed, the lattice can still spontaneously disassemble. In particular, at timescales less than the time it takes to reach this critical size, which even at 1.6μM adaptor is ^~^30s, lattices show a significant amount of dynamic remodeling. Most of the growth and shrinking occurs through monomers (Fig. S3B), and when a lattice does change size, the probabilities to dissociate a trimer or add one are comparable before reaching the maximal size limit (Fig. S3C). With changes to clathrin concentration, the size of initially stable nuclei does change, indicating that it is the total clathrin available that controls the initial barrier to nucleation, given sufficient adaptors (Fig. S4).

### Assembly in solution is less cooperative than on the membrane, despite increased stability of the curved structures

The above simulations tracked flat clathrin lattices on a flat surface. However, in solution cages are curved^51^, indicating that curved cages are more stable, absent the external constraints of the membrane. Recent experiments also quantified that clathrin assembly is more stable onto more highly curved surfaces, illustrating the energy gain that accompanies curved clathrin contacts^57^. Consistent with these observations, when we use the same energetic and kinetic parameters to simulate curved cage formation in solution, we see minimal assembly (Fig. 5). This shows that the total energy of the lattice on the membrane is weaker than that in solution, either because it is flat or because it is curved but its stability is lessened by forces from membrane bending. By increasing the binding free energy of the clathrin-clathrin contacts, consistent with increased stability of the curved lattice in solution, and by reducing the strain penalty on hexagons or pentagons, we see a growth in the assembly. This assembly yield is dependent on the adaptor concentration, which is consistent with *in vitro* experiments on adaptor-peptide driven cage formation in solution^51^, and shows significantly less cooperativity then on the membrane, due to the lack of dimensional reduction. By stabilizing the bonds at ^~^2k_B_T, and reducing the hexagonal strain by 2.3k_B_T, our cage assembly agrees relatively well with *in vitro* experiments (Fig. S5)^51^. As a result of both bond and hexagonal strain, we estimate that the curved lattice has an additional 3-4 k_B_T free energy per trimer (Table S1) available to perform work on the flat membrane, that is, to bend it. We refer to this quantity subsequently as the “clathrin curvature free energy”.

**Figure 5.**
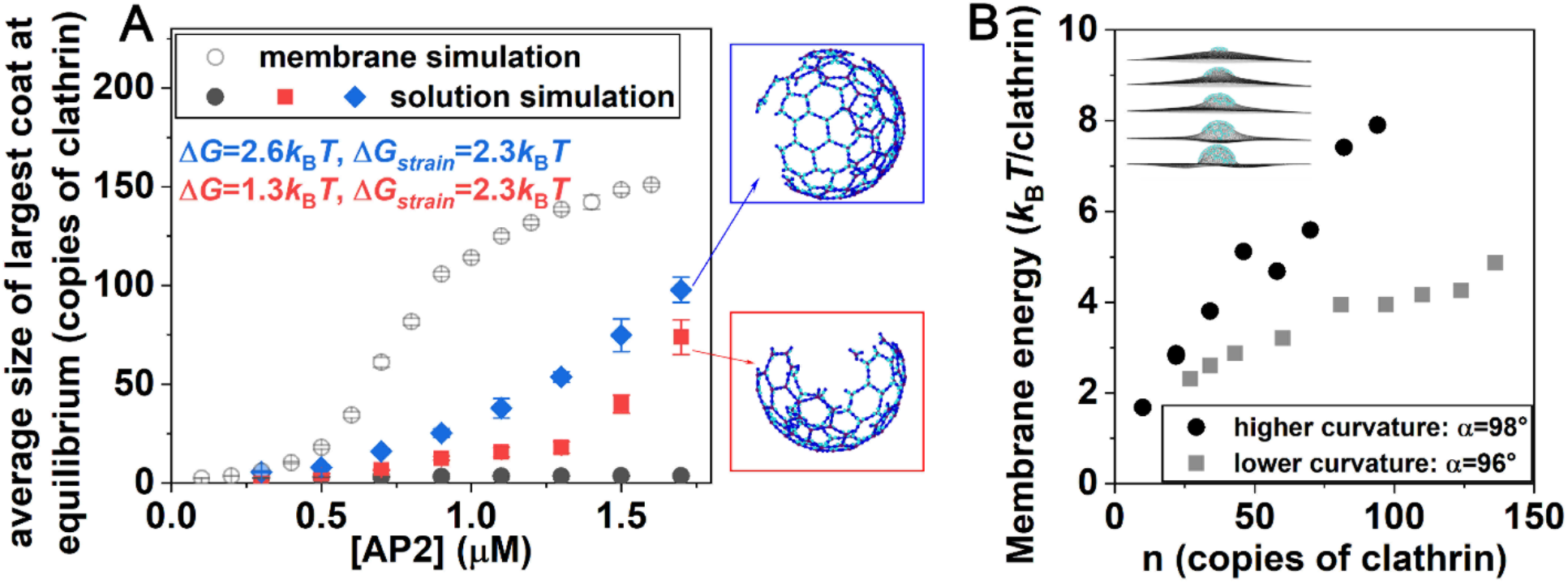
Curved-cage assembly in solution requires added stabilization, which is comparable to membrane bending energies. A) Clathrin cages with adaptors present do not form in solution (black dots), using same model as on the membrane (open circles). With added stabilization via clathrin-clathrin bonds (ΔG) and reduced strain (ΔG_strain_), solutions cages start to form (red and blue). B) Membrane bending energy per clathrin trimer as the sizes *n* of curved lattices are increased. More highly curved cages (*α*=98°) cost more energy per trimer to bend the membrane. The membrane energy is proportional to the bending modulus, where we use 20 k_B_T consistent with measurements on the plasma membrane^58^.

### Membrane bending cost can be at least partially compensated by stability of curved lattice

When formed on the membrane, curved cage formation must couple to membrane bending, and here we have harmonically linked our assembled curved cages with a deformable membrane model^59^ that follows the Helfrich Hamiltonian^60^ (Fig. 5B insets) (Methods). Membrane energies plotted in Fig. 5B indicate that the elastic energy of the membrane and the clathrin curvature free energy are comparable (3-4 k_B_T), indicating that curved pits are close in stability to the flat lattice. For the flat lattice on the membrane, it is the clathrin lattice that is strained, with the membrane at rest. When a pit forms, the strain on the lattice is eased, but the cost of the membrane bending increases. Note that we do not expect the curved, membrane bound clathrin lattice to recover the stability of the curved solution cages, due to forces placed on it by the deformed membrane. Thus curved assembly energies are modified on membranes, which was observed in a coarse-grained model of clathrin^43^. This stress balance is the basis for using separate rates for the curved lattices in solution vs expected values on the membrane, which would then be more similar to flat lattice values. Our calculations also demonstrate how the bending energy cost varies with the curvature of the vesicle. With larger vesicles, more clathrin are needed but the overall curvature is lower, resulting in a lower bending energy cost per clathrin (Fig. 5B).

### Discussion

Our model predicts that nascent clathrin lattices on membranes are unstable until ^~^25 clathrin are present, given physiologic parameters of 0.65μM clathrin and V/A=1μm. Our estimate of the stability threshold is remarkably similar to the sizes of abortive structures in cells^1^. A minimal requirement to overcome this barrier to nucleate structures on a membrane is a sufficient concentration of adaptor proteins that can link clathrin to the membrane, allowing assembly to cooperatively benefit from increased (adaptor-driven) stability and dimensional reduction. Physiological concentrations of AP-2 alone are insufficient to recruit enough clathrin to make membrane-bound cage formation spontaneous. Therefore, the participation of other adaptors is essential to drive nucleation. Our model shows how *in vitro* conditions with *in vivo* protein concentrations do not sample the transition between unstable and stable lattice growth, in large part due to an unphysiological V/A ratio. Our kinetic simulations and theoretical analysis allow us to bridge the geometric divide between *in vitro* and *in vivo;* our model predicts that for one adaptor type to drive productive nucleation and growth in ^~^60s (average time for vesicle formation) requires at least 1.6μM adaptors. This also requires sufficient lipid populations that can trigger efficient binding to the membrane, with slower lipid binding rates or reduced populations ultimately slowing both lag and growth rates. Our results clearly demonstrate that spontaneous disassembly of clathrin-coated structures on membranes can occur naturally even after relatively long time periods (many seconds). As evidenced by our equilibrium results, the accumulation of the majority of clathrin into single structures also depletes local clathrin and adaptor populations, leaving remaining clathrin capable of forming only small and transient structures (Fig. 4).

By modeling only a single type of adaptor for recruitment of clathrin to the membrane, we were able to construct predictive formulas for how the microscopic variables change the macroscopic lag and growth time, even as we varied the clathrin concentration, lipid concentration, system geometry, changes in cooperativity (Fig. S6) or rates of lipid binding (Fig. S7). However, *in vivo* there are multiple types of adaptors or accessory proteins that can link clathrin to the membrane^61^. Although our model predicts how adaptors with faster or slower binding kinetics would alter macroscopic growth, the ability of many of these proteins to bind to one another^62^ creates cross-linking that will stabilize these proteins on the membrane surface^15^. By increasing the local density of clathrin-recruitment sites, we predict this cross-linking would then help nucleate stable lattices at lower concentrations than occur for a single adaptor type. The binding kinetics of additional interactions can also be significantly more dynamic than AP-2; the proteins FCho1 and eps15 form clusters with AP-2 that helps initiate sites of clathrin-coated structure formation in cells, and yet both FCHo1 and eps15 are largely absent from completed vesicles, indicating the transience of their clathrin contacts^63–65^. The behavior of these additional adaptor protein types is consistent with our model prediction that AP-2 needs help to nucleate stable lattices, and in future work we will consider how explicit cross-linking can enhance local concentrations to drive nucleation and growth in time and space. Importantly, the model and software are open source, so they can be immediately applied and modified with additional experimental data (Methods).

A limitation of our model is the clathrin-coat assembly is not dynamically coupled to the membrane remodeling to form spherical vesicles. However, our energetic calculations using our assembled clathrin structures and a deformable membrane model demonstrate that the energy per clathrin needed to bend the membrane is comparable to the clathrin curvature free energy. Our model thus predicts that the energy of flat lattices on membranes and curved cages on bent membranes is of a similar scale, albeit sensitive to the membrane stiffness and additional curvature generation mechanisms. Additional curvature induction driven by adaptor proteins (e.g. amphipathic helix insertion and BAR domains^66^) would help remodel the membrane, thus reducing the work required from the clathrin lattice alone. The kinetics of membrane bending is unlikely to be rate-limiting, given the slow time-scales of clathrin recruitment and relatively fast remodeling dynamics observed in membrane budding^67^. The kinetics of late cage assembly could be altered due to budding, however, due to the reduction in clathrin lattice perimeter in a curved cage as the size proceeds beyond the vesicle diameter (^~^150nm). While this would not impact the early growth to overcome the nucleation barrier, the closing of the vesicle involves a shrinking perimeter that reduces sites to recruit clathrin, but increases contacts formed on average per clathrin. Modeling the kinetics of coupled assembly and remodeling will ultimately be important to isolate productive vesicle forming events, and is feasible for the types of models used here.

The clathrin coat assembly studied here shares many fundamental properties with diverse biological pathways including steps in viral assembly^67–-69^ and protein aggregation^70^, where we would expect similar sensitivity to membrane or surface (air/water) localization and geometry. Our work here predicts how assembly and aggregation can occur on surfaces with weaker interaction energies or significantly lower copy numbers than is needed to nucleate structures in solution. Localization to the surface provides an additional time-scale that can slow nucleation, which would be helpful in avoiding kinetic traps^71^ to improve yield in designed systems^72,73^. The addition of the membrane provides a control variable for experimental quantification of interaction kinetics, where we showed here how visualization of recruitment to surfaces^25^ can be an effective observable for discriminating assembly mechanisms. Enzymatic control of membrane composition can then be used to trigger disassembly, to study unbinding rates^28^. Overall, the rate-based approach used here^28^ offers a valuable platform for mechanistic and predictive modeling of self-assembly, in and out-of-equilibrium.

## METHODS

### Reaction-diffusion simulations

Simulations are performed using the NERDSS software^28^, which propagates particle-based reaction-diffusion using the free-propagator reweighting (FPR) algorithm^30^. The lipids or adaptors restricted to the membrane are modeled using an implicit lipid algorithm, that reproduces the same kinetics and equilibria as the more expensive explicit lipid method^32^. We use a time-step Δt=3μs. The accuracy and reproducibility of the simulation results is verified using a smaller time-step (Fig. S1B). Software and executable input files for the models used here are provided open source at github.com/mjohn218/NERDSS.

### Continuum membrane calculations

The membrane is represented by a curvature-continuous spline via the subdivision limit surface algorithm as in Ref.^59^. The surface, formed by approximately hexagonal triangles, uses periodic boundary conditions with side length 700 nm, for an area of 490000 nm^2^. Energetically the membrane is restrained to match the cage according to harmonic potentials, one for each clathrin. A virtual site was placed approximately 9 nm from the clathrin center of mass (ca. 7.5 nm from the adaptor sites). A distance 2.5 nm closer was also attempted with comparable energetics (within 2%). The distance between the virtual site and the membrane patch was determined as in Ref.^32^. This nearest point on the membrane, tagged in local surface coordinates (face, and internal triangular coordinates) was saved during 50 “sub-iterations” of the Broyden-Fletcher-Goldfarb-Shanno (BFGS) conjugate-gradient class algorithm. A minimization schedule was found that yielded robust fits. Force constants were gradually increased by a factor of three, beginning with 0.0003 and stopping with 100 kcal/mol/Angstrom^2^. The clathrin cage was allowed to shift and rotate during minimization. The clathrin cages were generated via NERDSS simulations in solution, using an angle α=98⍛ (see Fig 1) and a softer curvature of 96⍛.

### Concentrations

The values for clathrin and AP-2 copy numbers in HeLa cells, along with the cell sizes, are taken from this paper^62^, compiled from experimental studies^56^. For AP-2, we use the concentration of the limiting subunit, and for clathrin trimers, we use copies of clathrin heavy chain divided by 3.

### Experimental system size

The experimental V was 200μL^25^ and the experimental A of 2.017×10^8^ μm^2^ is calculated based on 1 nmol of total lipid, with an average lipid SA of 0.7nm^2^ and a two-leaflet bilayer forming cylindrical tubules. The experimental V/A ratio is then 991 μm.

### Estimating n_1_ and n_2_ lattices sizes from probability distributions

To extract the lattice sizes where the barrier to initial growth flattened (Fig. 4), we analyzed the -ln(P(n)) distribution for each trajectory. We chose a plateau value in -ln(P(n)) for each trajectory, and where this value was first crossed determined n1, and where it was last crossed (at the descent into the stable well) determined n_2_ (Fig. 4B). The plateau value was chosen based on where -ln(P(n)) transitioned to noisy fluctuations, where we note that choosing a higher value would predict slightly larger values of n_1_ and smaller values of n_2_. For the equilibrium distributions, there was typically a gap where no lattices of intermediate sizes were observed. The start and end of this gap were used to determine n_1_ and n_2_.

## ACKNOWLEDGEMENTS

M.E.J. gratefully acknowledges funding from a National Institutes of Health MIRA Award R35GM133644. A.J.S. is supported by the Intramural Research Program of the NIH. We acknowledge use of the ARCH supercomputers Bluecrab (MARCC) and Rockfish at Johns Hopkins, and the XSEDE supercomputer Stampede2 through XRAC MCB150059. We are grateful to Prof Thomas Pucadyil for sharing his data and for helpful discussions on the experiments.

## Supplementary Information Text

### SI METHODS

#### Maintaining a constant (fluctuating) clathrin concentration

The constant clathrin concentration is maintained by a zeroth order reaction producing clathrin at *k*_create_=[Cla]_tot_D_Cla_/z^2^, which approximates the time-scales of diffusive flux across volume elements of height z. Simultaneously, clathrin is degraded in solution at a rate of *k*_destroy_=D_Cla_/z^2^, where D_Cla_ is set to 13μm^2^/s. Clathrin on the membrane is not affected by the degradation reaction.

#### System parameters of the NERDSS simulation for assembly reactions

Association events within a rigid complex are necessary to close polygons (like hexagons and pentagons). These unimolecular reactions are attempted when reactive partner sites are within a maximum distance defined by the binding radius σ multiplied by a factor, bindRadSameCom ^74^. The bindRadSameCom was set to 1.1 in our simulations.

NERDSS can reject associate events for two reasons:

1. if an association event causes components of the two complexes to sterically overlap, the event is rejected, and the complexes remain separate. NERDSS evaluates steric overlap after association by measuring distances between all centers of mass (COMs) in the new complex and defining a minimum distance, overlapSepLimit, that will determine the steric overlap^74^. The overlapSepLimit was set to 7nm in our simulations.
2. if an association event causes elements of either complex to undergo large-scale displacement as they snap into the proper orientation, the move is rejected. This effectively only applies to large complexes, where sites distant from the COM can potentially displace significantly upon association. This is enforced by a cutoff value, where events that result in shifts of an interface on either complex by scaleMaxDisplace* <RMSD> are rejected, because the displacements are not physical. <RMSD> is calculated from (6*D_eff_*Δt)^0.5^ in 3D, and (4*D_eff_*Δt)^0.5^ in 2D, where D_eff_ is the effective translational diffusion coefficient of one complex ^29^, accounting for rotational diffusion ^74^. The scaleMaxDisplace was set to 10 in our simulations.

#### Mesh resolution in the continuum membrane model

Mesh resolutions were tested from 448 to 1344 triangles (ca. 1093-364 nm^2^ per triangle) with variation in the energy with mesh resolution on the order of 5% for highly deformed structures. The energy generally decreases with increased mesh resolution as it has more freedom to adapt to the cage. The energy is determined as a function of the mesh vertex coordinates that are input to the subdivision scheme.

During minimization, the nearest point on the membrane, tagged in local surface coordinates (face, and internal triangular coordinates) was saved during 50 “sub-iterations” of the Broyden-Fletcher-Goldfarb-Shanno (BFGS) conjugate-gradient class algorithm. Following these sub-iterations, new nearest points were computed for each virtual site. In this way, efficient BFGS minimization could proceed without the somewhat expensive recalculation of the nearest sites.

#### Transition matrix and probability distribution of clusters with different size

NERDSS tracks all complexes formed based on the number and type of components. We calculate the transition matrix T(Δt) using a small time interval for high resolution, Δt=150μs, to measure growth probabilities and lifetimes (Fig. S3). The diagonal of T(Δt), T(Δt)_*n,n*_ contains the probability that one cluster stays the same size n in a time step Δt. Off the diagonal are transitions between clusters of all sizes, from *n*=1 to *n*=N_max_ (total number of clathrin) including monomer and non-monomer transitions. P(*n*) is the probabilities of observing a cluster with size *n* and can be calculated from summing T(Δt) over the column *n*, and then normalizing over all *n*.

#### Phenomenological fit to lag and initial growth rate

##### Lag time

The lag time was fit to a sum of timescales, each of the form of (*rate* x *concentration*)^-1^. To describe the full set of simulations, a third timescale was added to account for binding of the adaptor proteins to the lipids. This term was not needed to describe the *in vitro* data, as adaptors were already localized to the surface (it was zero).

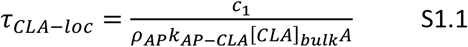

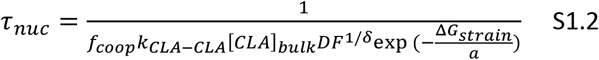

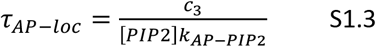

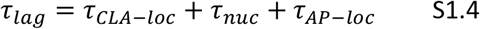

The fit parameters were found using nonlinear fitting in MATLAB, against lag times collected across 80 different simulation points (Table S2). The correlation between the observed lag times and the model predictions using Eq S1.4 are in Fig. S2. Optimal fit values are printed in Eq 2, and *c*_3_ = 0.4.

##### Steepness/Initial growth rate

For growth, we used a mass-action type expression, which similarly had three terms for the full system, with the third term only needed when binding to lipids occurred.

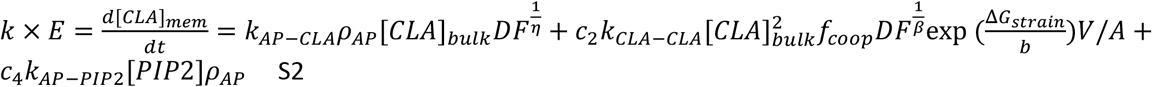

Fit parameters were again found using nonlinear fitting in MATLAB against 80 different simulations points, with correlations shown in Fig. S2. This function fails to fully capture the dependence of the growth rate on cooperative clathrin-clathrin growth, systematically underpredicting growth rates with higher clathrin concentrations. Optimal fit values printed in Eq 3, and *c*_4_ = 0.00078. The steepness is measured in copies/s/μm^2^. We note that the dimensionless *c*_2_ value is fit using concentrations in μM, V in units of μm^3^, and A in units of μm^2^. Cancellation between the units of μm^3^ and μM is a factor of 602, which is included in *c*_2_.

#### Energies per trimer in clathrin lattices

To calculate average energies per trimer in a lattice, we use approximate expressions derived assuming the clathrin grow into a compact lattice, maximizing formation of hexamers ^29^. We thus use:

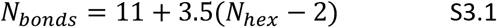

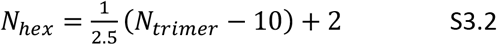

## Supplemental Table

**Table S1.** Calculations of average energies per trimers assembled into lattices, due to bonds formed and hexamers formed. Counts from Eq S3.

| Counts per lattice |  |  | Energy per bond: -11.7kBT, Energy strain per hexamer: 6.9kBT |  | Energy per bond: -14kBT, Energy strain per hexamer: 4.6kBT |  |  |
| --- | --- | --- | --- | --- | --- | --- | --- |
| N trimers | N hexamer | N bonds | Total energy on membrane (kBT) | Energy per trimer (kBT) | Total energy in solution (kBT) | Energy per trimer solution (kBT) | Total added stability per trimer (kBT) |
| 5 | 0 | 4 | -46.8 | -9.36 | -56 | -11.2 | -1.84 |
| 10 | 2 | 11 | -114.9 | -11.49 | -144.8 | -14.48 | -2.99 |
| 20 | 6 | 25 | -251.1 | -12.555 | -322.4 | -16.12 | -3.565 |
| 30 | 10 | 39 | -387.3 | -12.91 | -500 | -16.666667 | -3.7566667 |
| 50 | 18 | 67 | -659.7 | -13.194 | -855.2 | -17.104 | -3.91 |
| 100 | 38 | 137 | -1340.7 | -13.407 | -1743.2 | -17.432 | -4.025 |
| 150 | 58 | 207 | -2021.7 | -13.478 | -2631.2 | -17.541333 | -4.0633333 |
| 200 | 78 | 277 | -2702.7 | -13.5135 | -3519.2 | -17.596 | -4.0825 |

**Fig. S1.**
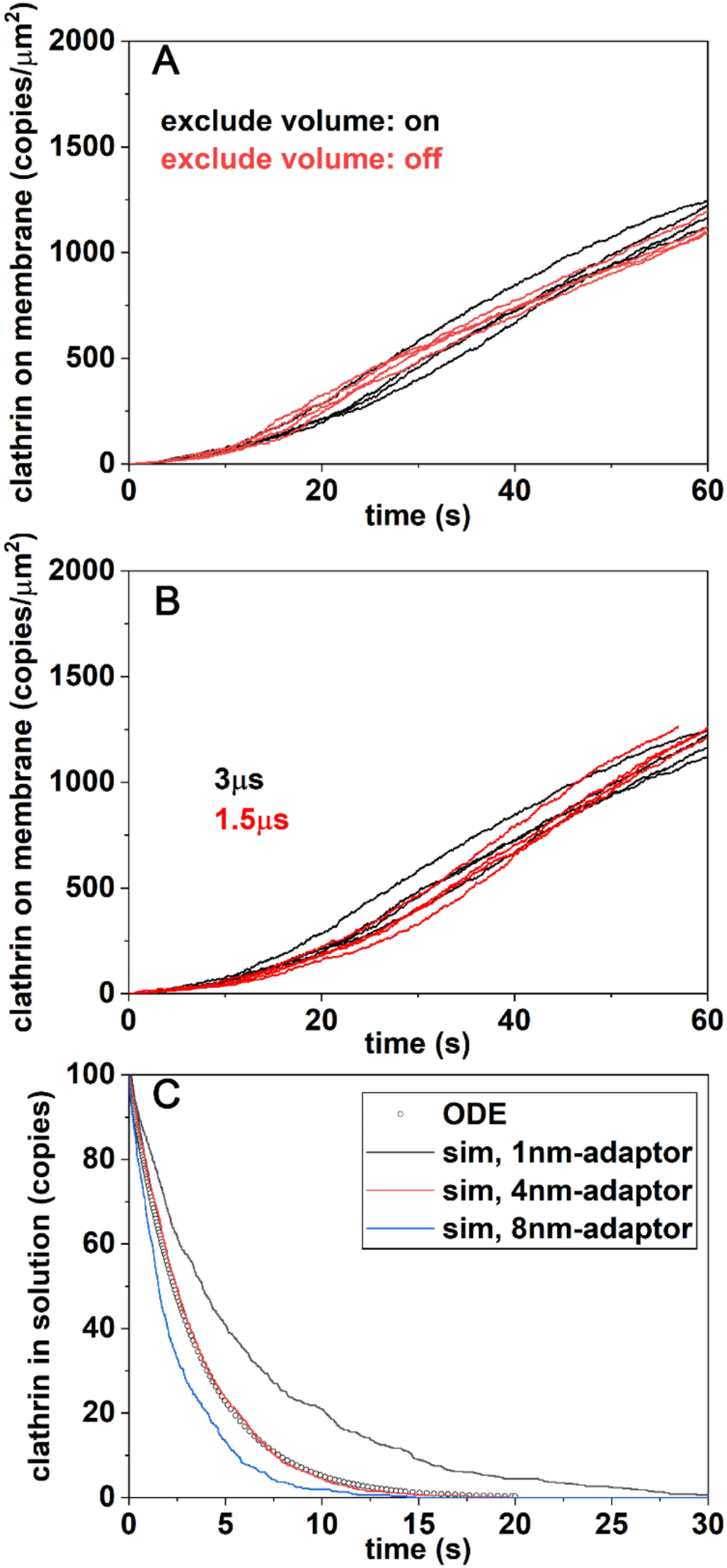
Numerical confirmation of expected kinetics. A) Excluded volume between clathrin centers does not affect the kinetics. B) The kinetics is independent of the simulation time-step. C) The adaptor with a height of 4nm ensures accurate membrane binding kinetics. This separation of the binding site above the reflective membrane surface is necessary due to the extended size of the rigid multi-site clathrin. The reflection of all the clathrin sites off of the membrane surface can reduce reactive flux between the reactive binding sites of clathrin-adaptor. The system box is [1000, 1000, 1000] nm, including 100 copies of clathrin, 50000 copies of implicit adaptors on membranes, and the binary reaction between clathrin and adaptor. k_on,3D→2D_=0.0012μM^-1^s^-1^. k_on,2D_=0.04μM^-1^s^-1^. k_off_=0.03s^-1^. The ODE is calculated using the VCell^75,76^ ignoring the spatial structure of molecules. The sim is 6 trajectories generated from simulation of NERDSS^74^.

**Fig. S2.**
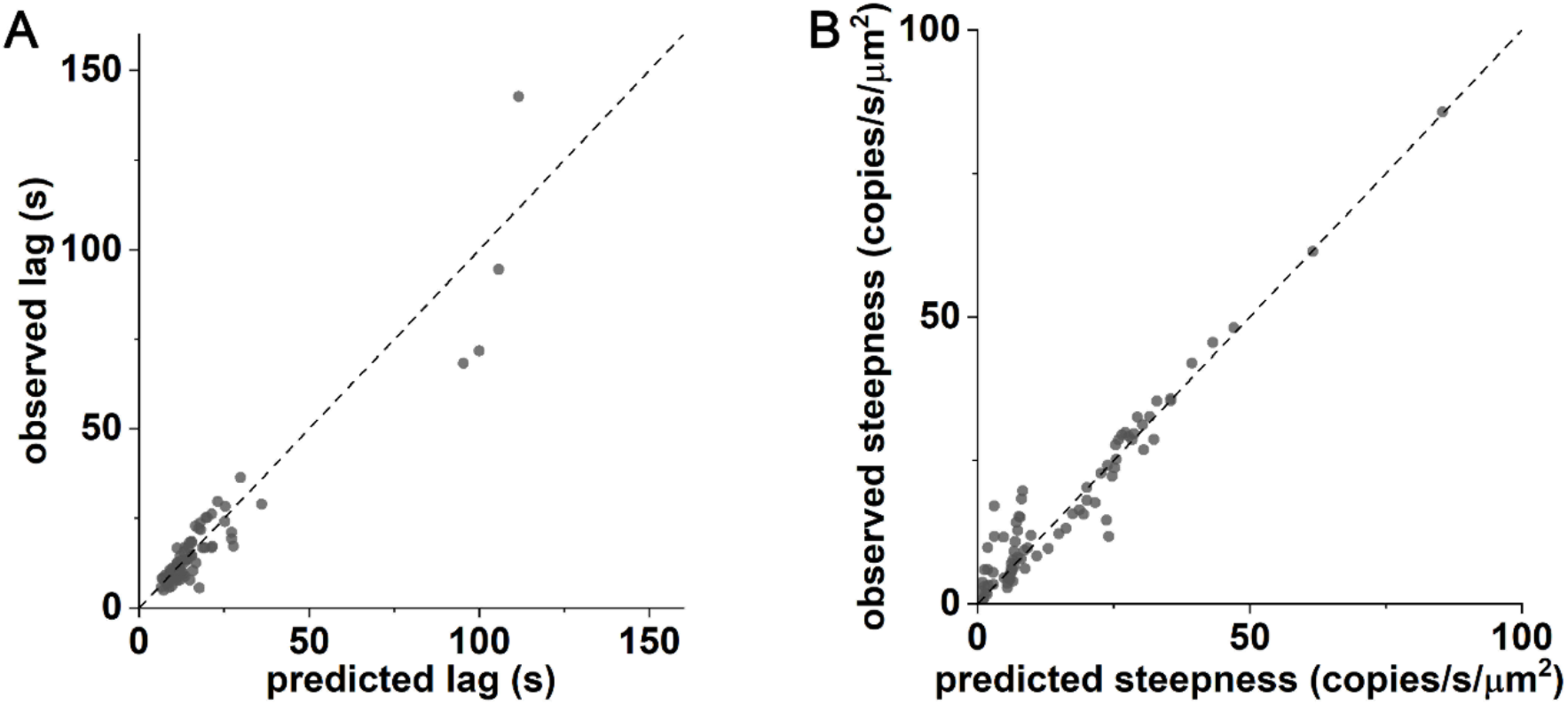
Correlations between observed lag and initial growth rate vs predicted values from the simplified models of Eq 2 and Eq 3. A) Predicted lag agrees quite well with observed lag, with results shown for 80 simulations (Table S2). B) Predicted initial growth rate, or steepness, agree relatively well with observed values. Same 80 simulations.

**Fig. S3.**
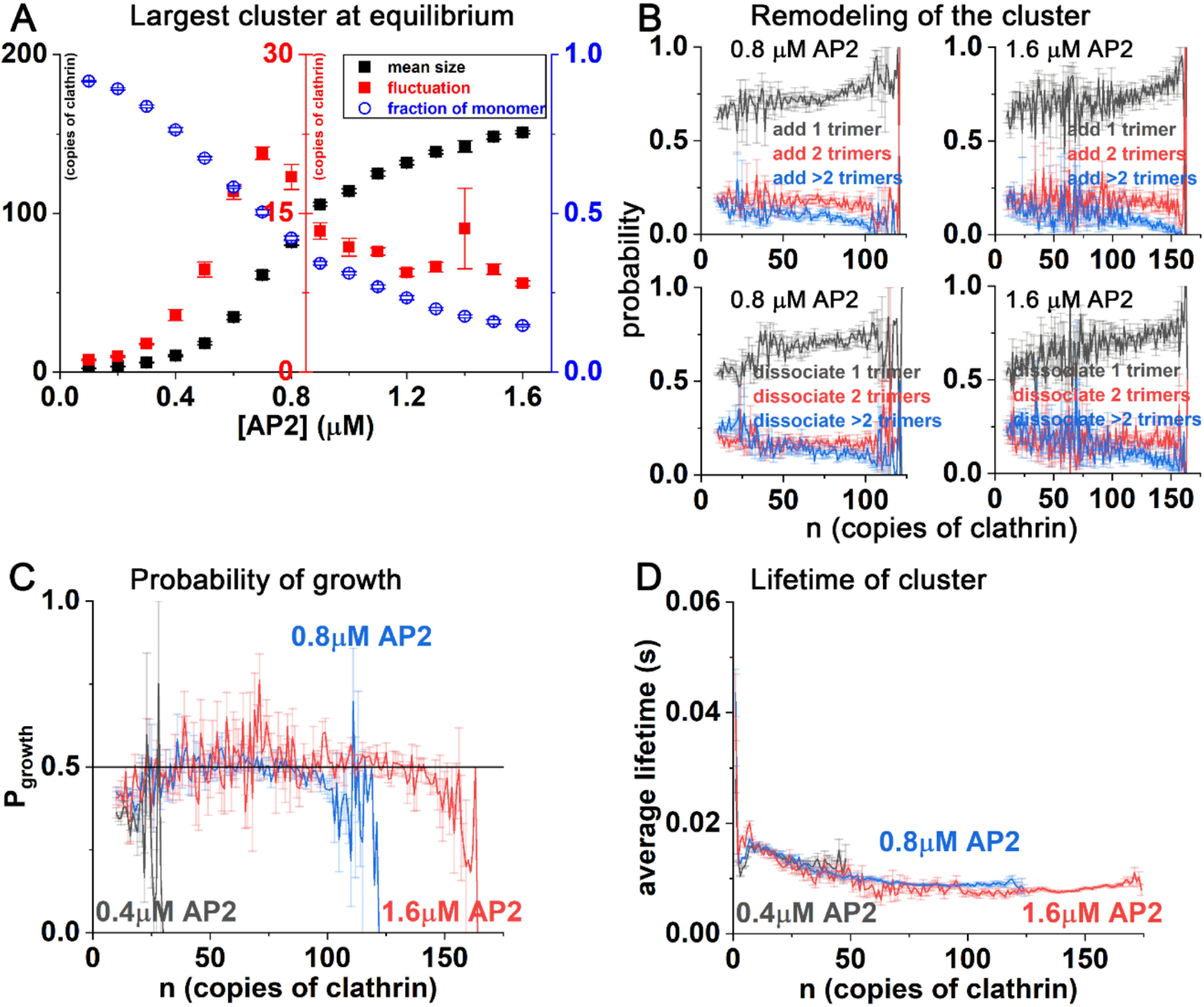
Growth mechanisms of clathrin lattices at the more “physiologic-like” geometries. A) Sizes of clathrin structures vs adaptor concentration. All data points have the same clathrin concentration (0.65μM). The mean lattice size increases with adaptors (black), with a corresponding drop in the fraction of monomers (blue). The fluctuations in the lattice sizes at equilibrium peaks at 0.7μM (red) indicating a transition point separating small and large stable lattices. B) The growth of lattices of size n (x-axis) typically proceeds by addition or removal of a monomer (black data). About 20% of the growth is due to addition or removal of dimers (red), and the remaining is due to higher-order structures (blue). C) Given that a transition occurs from a lattice of size *n* (x-axis) to a new size, we plot the probability that the transition causes a growth in size. For small lattices, the growth probability is less than 0.5, meaning that disassembly is more probable. For intermediate sizes, there is a small bias towards growth, until very large lattices form. This supports the dynamic remodeling observed in the simulations, as at all sizes, there is still a significant probability of disassembly. D) The lifetimes of lattices of size *n* is ^~^0.01-0.02s, with monomers being the longest lived.

**Fig. S4.**
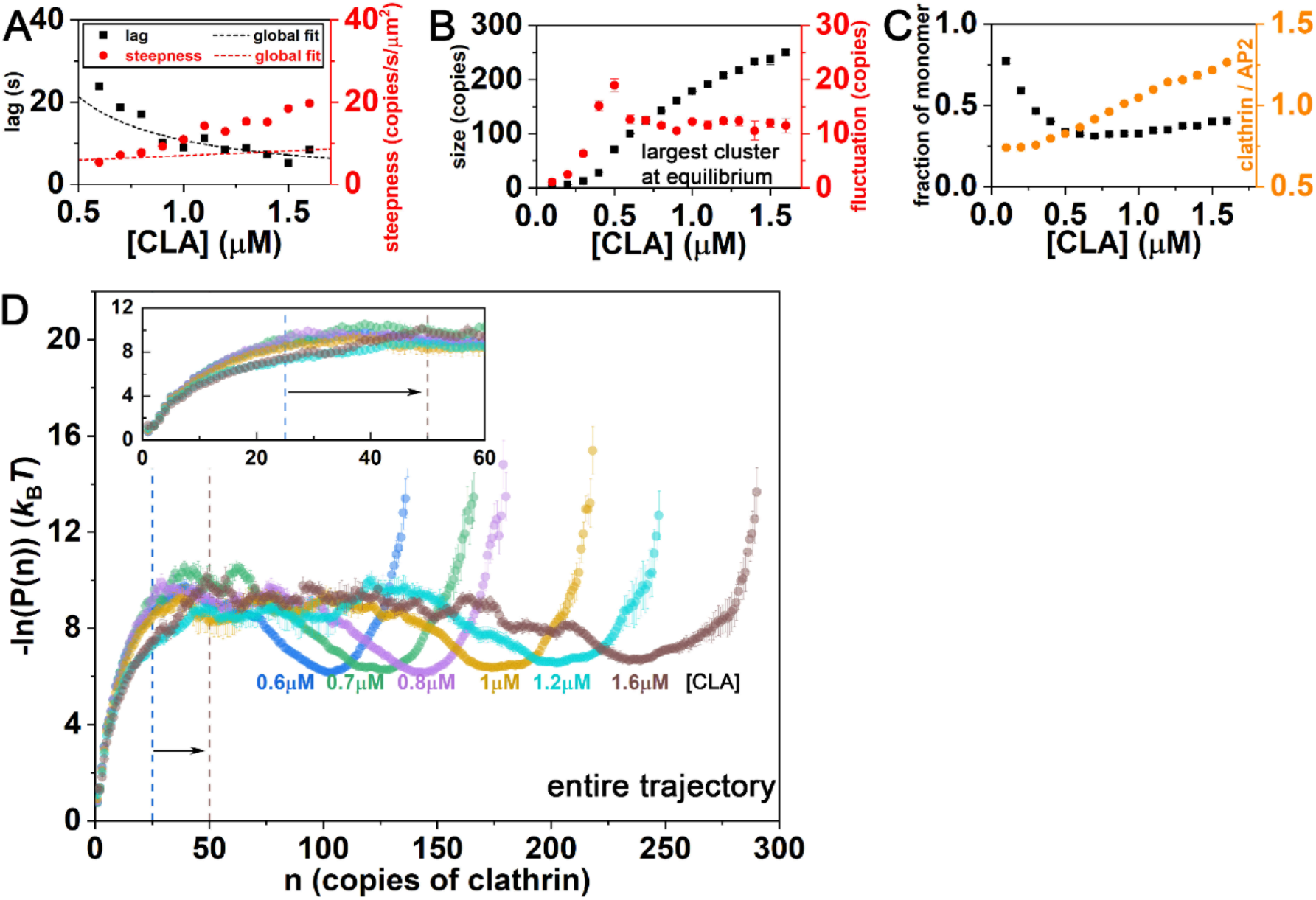
Growth of lattices with changing clathrin concentration, at fixed 1μM of adaptors. A) Lag time is well-described by the main text Eq 2/Eq S1. Growth rate of Eq 3/Eq S2 does as accurately capture speed-ups with higher clathrin concentration. The simple formula does not have sufficient cooperativity in clathrin-clathrin contacts in stimulating growth. B) Lattice sizes increase with increasing clathrin concentration (black) with fluctuations peaking at 0.5μM clathrin. Prior to this point, maximal clathrin lattices are quite small C) The fraction of monomeric clathrin decreases before flattening out (black). The stoichiometry of clathrin bound to adaptors keeps increasing, as more clathrin is added. It is not sufficient to have a stoichiometry of at least 1:1; at 0.2μM clathrin, the stoichiometry is more like 1:1.5, and yet no lattices form. Thus the total concentration is also a significant determinant of nucleation and growth. D) The probability of observed cluster sizes *n* shifts to larger lattices with additional clathrin. The initial barrier also appears to shift to larger sizes with additional clathrin.

**Fig. S5.**
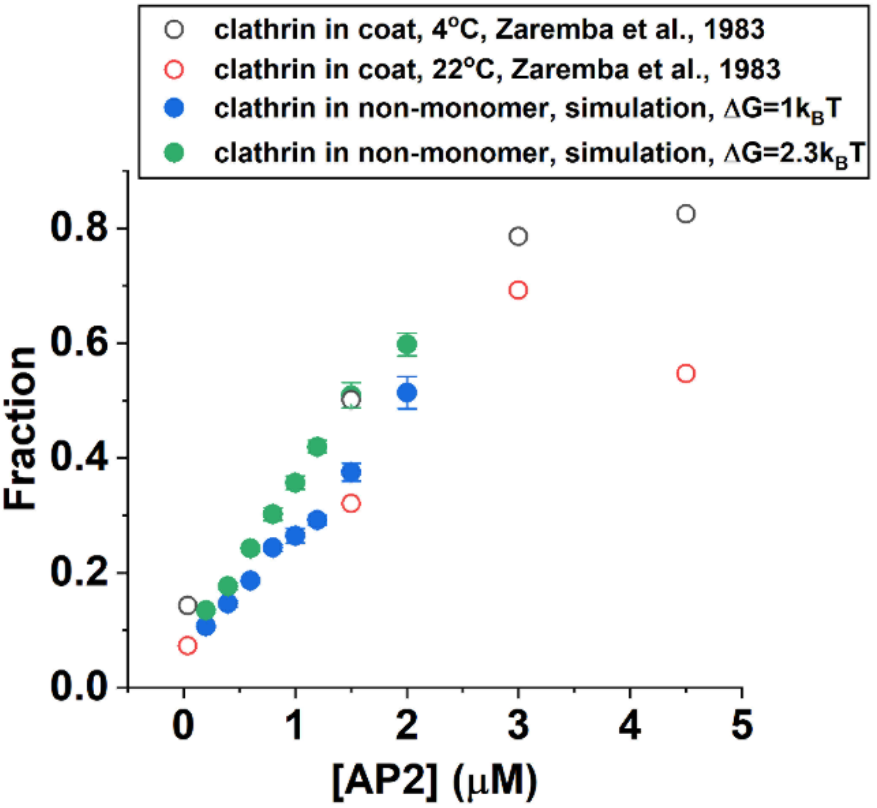
Fraction of assembled clathrin in solution vs adaptor concentration. Experimental data is (open red and black circles) from Ref ^51^. To convert from mg/ml of protein to μM, we used a molecular mass of ^~^650kDa for clathrin, and 50kDa for the adaptors, based on the values reported in Table 1 of Ref ^51^. For the simulations, the clathrin concentration is 0.65μM. The clathrin concentration in our simulations is thus higher than in the experiments (^~^0.3μM), meaning that our model does not cause as strong assembly as observed experimentally. We note that in the experiments, the adaptors represented a mixture, that easily could have clustered amongst themselves, helping to nucleate lattices via crosslinking. Thus, we overall do not consider quantitative reproduction of these older experiments as critical as the recent fluorescence experiments.

**Fig. S6.**
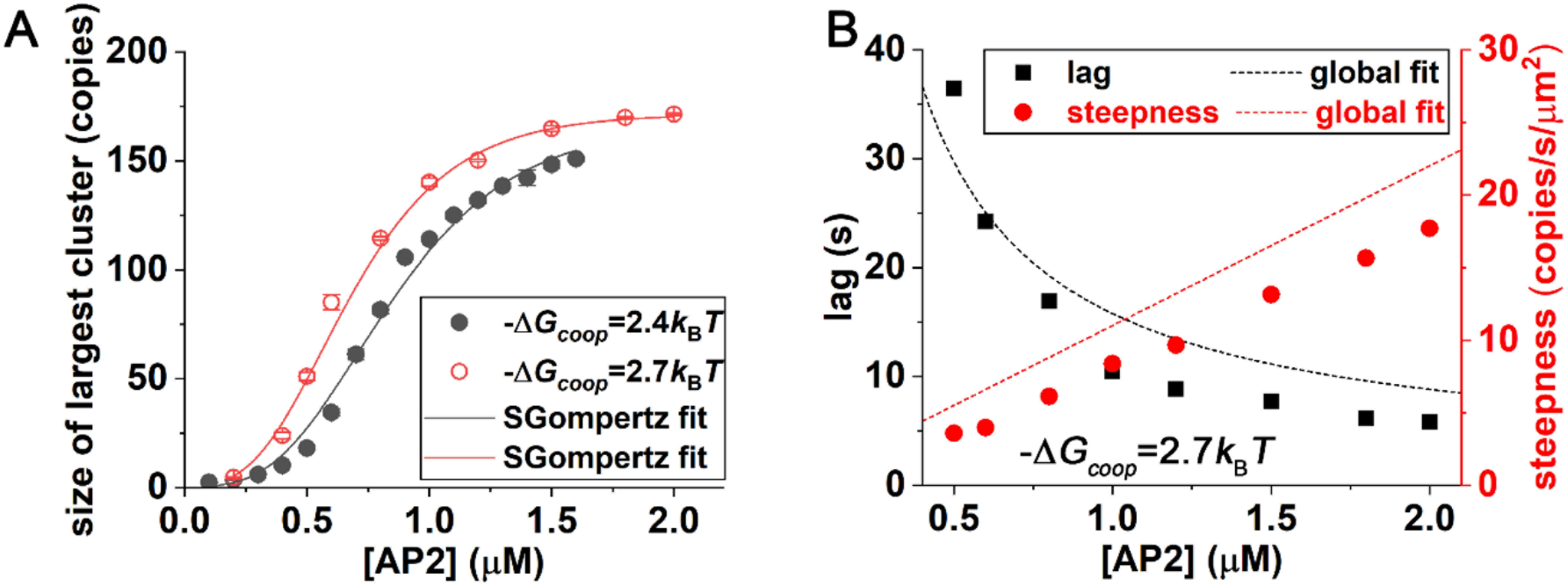
Maximal lattices formed as AP2 concentration is varied, comparing two values of cooperativity in clathrin-clathrin contacts, ΔG_coop_. A) For adaptors that provoke stronger cooperativity in clathrin-clathrin assembly (red), the overall trend is the same, but the transition to larger and more stable lattices occurs at lower adaptor concentration. B) For the more cooperative model, the lag-time and initial growth rate are described well by Eq 2 and Eq 3.

**Fig. S7.**
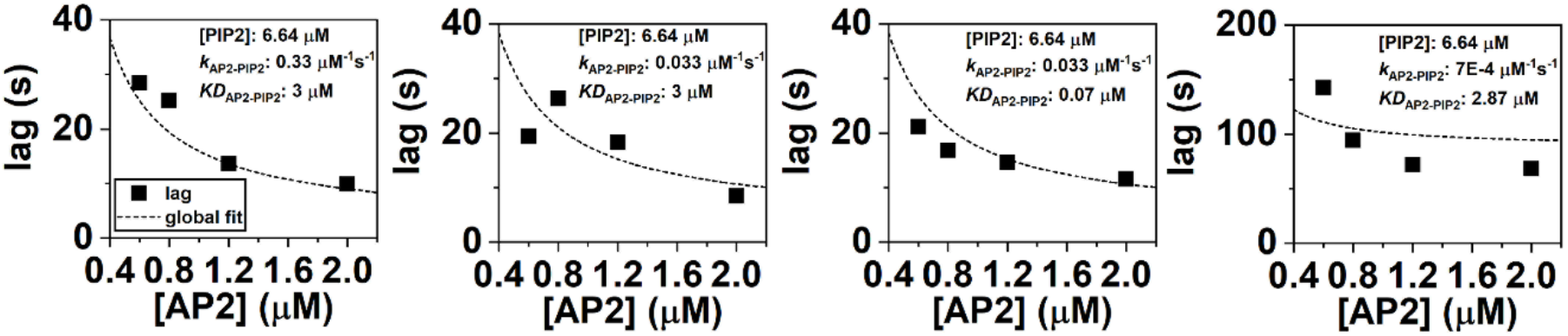
Reduced lipid populations or slower lipid binding rates slow the lag. The lag-time are described well by Eq S1 as the lipid concentration, or rate and strength of adaptor-lipid binding are varied.

---

**Movie S1 (separate file).** Video of the clathrin assembly matching the *in vitro* simulations, as shown in Fig 2.

**Table S2 (separate file).** Excel sheet containing the parameters for all simulations, along with observed lag times and growth rates, and predicted lag times and growth rates.

